# LHP1 and INO80 cooperate with ethylene signaling for warm ambient temperature response by activating specific bivalent genes

**DOI:** 10.1101/2024.03.01.583049

**Authors:** Zhengyao Shao, Yanan Bai, Enamul Huq, Hong Qiao

**Affiliations:** Institute for Cellular and Molecular Biology, The University of Texas at Austin, Austin, TX, 78712, USA; Department of Molecular Biosciences, The University of Texas at Austin, Austin, TX, 78712, USA

## Abstract

Ethylene signaling has been indicated as a potential positive regulator of plant warm ambient temperature response but its underlying molecular mechanisms are largely unknown. Here, we show that LHP1 and INO80 cooperate with ethylene signaling for warm ambient temperature response by activating specific bivalent genes. We found that the presence of warm ambient temperature activates ethylene signaling through EIN2 and EIN3, leading to an interaction between LHP1 and accumulated EIN2-C to co-regulate a subset of LHP1-bound genes marked by H3K27me3 and H3K4me3 bivalency. Furthermore, we demonstrate that INO80 is recruited to bivalent genes by interacting with EIN2-C and EIN3, promoting H3K4me3 enrichment and facilitating transcriptional activation in response to warm ambient temperature. Together, our findings illustrate a novel mechanism wherein ethylene signaling orchestrates LHP1 and INO80 to regulate warm ambient temperature response through activating specific bivalent genes in Arabidopsis.

## Main

Over the past few decades, global warming resulting in gradual increase in temperature has placed significant challenges on plant growth, development, and more importantly crop yield^1, 2^. Ethylene signaling has been indicated as a potential positive regulator of warm ambient temperature^3^. Mutations in ethylene receptor *ETHYLENE RESPONSE 1 (ETR1)* and key factors in ethylene signal transduction pathway *ETHYLENE INSENSITVE 2 (EIN2)* and *ETHYLENE INSENSITVE 3 (EIN3)* result in a decreased thermotolerance; in contrast, a constitutive ethylene responsive mutant of *CONSTITUTIVE TRIPLE RESPONSE 1 (CTR1)* is more tolerant to heat shock treatment^4,5^. *HEAT SHOCK TRANSCRIPTION FACTOR A7 (HSFA7) HSFA7a* and *HSFA7b* are reported to regulate both ethylene signaling and ethylene biosynthesis homeostasis to properly establish thermotolerance and thermopriming at shoot apical meristem tissues^6^. Moreover, a recent study showed that EIN3 protein is stabilized by warm ambient temperature (27°C) due to the degradation of its negative regulators EIN3-Binding F box protein 1 (EBF1) and EBF2 by Salt- and Drought-Induced Ring finger 1 (SDIR1)^7^. These studies strongly suggest that ethylene signaling potentially functions as a positive regulator of warm ambient temperature. However, the molecular mechanisms underlying the involvement of the ethylene signaling in the warm ambient temperature response largely remain to be investigated.

During our study of ethylene and temperature responses, we found that the warm ambient temperature (27°C)-induced elongation in hypocotyl in Col-0 was reduced in the key ethylene signaling component mutants *ein2-5* and *ein3-1eil1-1* (Fig. S1A and S1B). The reduction was also observed in *lhp1-3,* the null mutant of a Polycomb repressive complex (PRC) component *LIKE HETEROCHROMATIN PROTEIN 1 (LHP1)* (Fig. S1A and S1B)^8^. To further investigate how ethylene and LHP1 are involved in warm ambient temperature response, we conducted histochemical staining for GUS activity of ethylene reporter *5xEBS::GUS* in response to warm ambient temperature^9^. The result showed that warm ambient temperature leads to an elevation of ethylene signaling (Fig. S1C). We then examined EIN2, EIN3, as well as LHP1 protein levels under a warm ambient temperature (27°C) treatment. EIN2 cleavage and EIN3 protein accumulation were induced by warm ambient temperature (Fig. S1D and S1E). However, the level of LHP1 protein remains unchanged (Fig. S1F).

Next, we compared the transcriptomes of Col-0, *ein2-5, ein3-1eil1-1*, and *lhp1-3* with or without four hours of 27°C treatment (Fig. S1G and S1H). We found that the warm ambient temperature-induced transcriptional activation in Col-0 was compromised in the *ein2-5, ein3-1eil1-1*, and *lhp1-3* mutants with a similar expression pattern (Fig. S1G and S1H). Subsequent statistical analysis further confirmed the reduction was significant in all three mutants compared to Col-0 (Fig. S1H). More importantly, the reductions observed in *ein2-5, ein3-1eil1-1*, and *lhp1-3* were statistically comparable, suggesting the potential co-functionality of EIN2, EIN3, and LHP1 in response to a warm ambient temperature.

To investigate the relationship between LPH1 and EIN2 or EIN3 in the context of a warm ambient temperature response, we first generated *ein2-5lhp1-3* and *ein2-5 lhp1-6* double mutants, and *ein3-1eil1-1lhp1-3* and *ein3-1eil1-1lhp1-6* triple mutants. Warm ambient temperature response assay showed a significant reduction in warm ambient temperature induced hypocotyl elongation in *ein2-5lhp1-3* and *ein2-5lhp1-6* double mutants as well as *ein3-1eil1-1lhp1-3* and *ein3-1eil1-1lhp1-6* triple mutants compared to the *ein2-5* and *lhp1-3* or the *ein3-1eil1-1* mutants, respectively (Fig. 1A and 1B). Notably, these higher order mutants exhibited almost no response to warm ambient temperature (Fig. 1A and 1B). Our further test with PHYTOCHROME INTERCTING FACTOR 4 (PIF4), the master regulator of thermomorphogenesis, protein levels showed that in *ein2-5, ein3-1eil1-1, lhp1-3,* as well as in *ein2-5lhp1-3* and *ein3-1eil1-1lhp1-3,* PIF4 protein is accumulated to the same level as in Col-0 under warm ambient temperature treatment, suggesting that the initial warm ambient temperature response by PIF4 accumulation is not impaired in these single mutants nor in higher order mutants (Fig. S2A and S2B)^10, 11^. To further validate the synergistic genetic function of LHP1 with EIN2 and EIN3 in warm ambient temperature ambient response, we introduced *proLHP1::gLHP1-GFP* into *lhp1-6*, *ein2-5lhp1-6*, and *ein3-1eil1-1lhp1-6* mutants and examined their responses to warm ambient temperature^12^. The phenotypes of *lhp1-6*, *ein2-5lhp1-6*, and *ein3-1eil1-1lhp1-6* were restored by *proLHP1::gLHP1-GFP* to that of Col-0, *ein2-5*, or *ein3-1eil1-1*, respectively (Fig. 1A and 1B and S2C), further confirming that LHP1 functions synergistically with EIN2 and EIN3/EIL1 in responses to warm ambient temperature.

**Figure 1.**
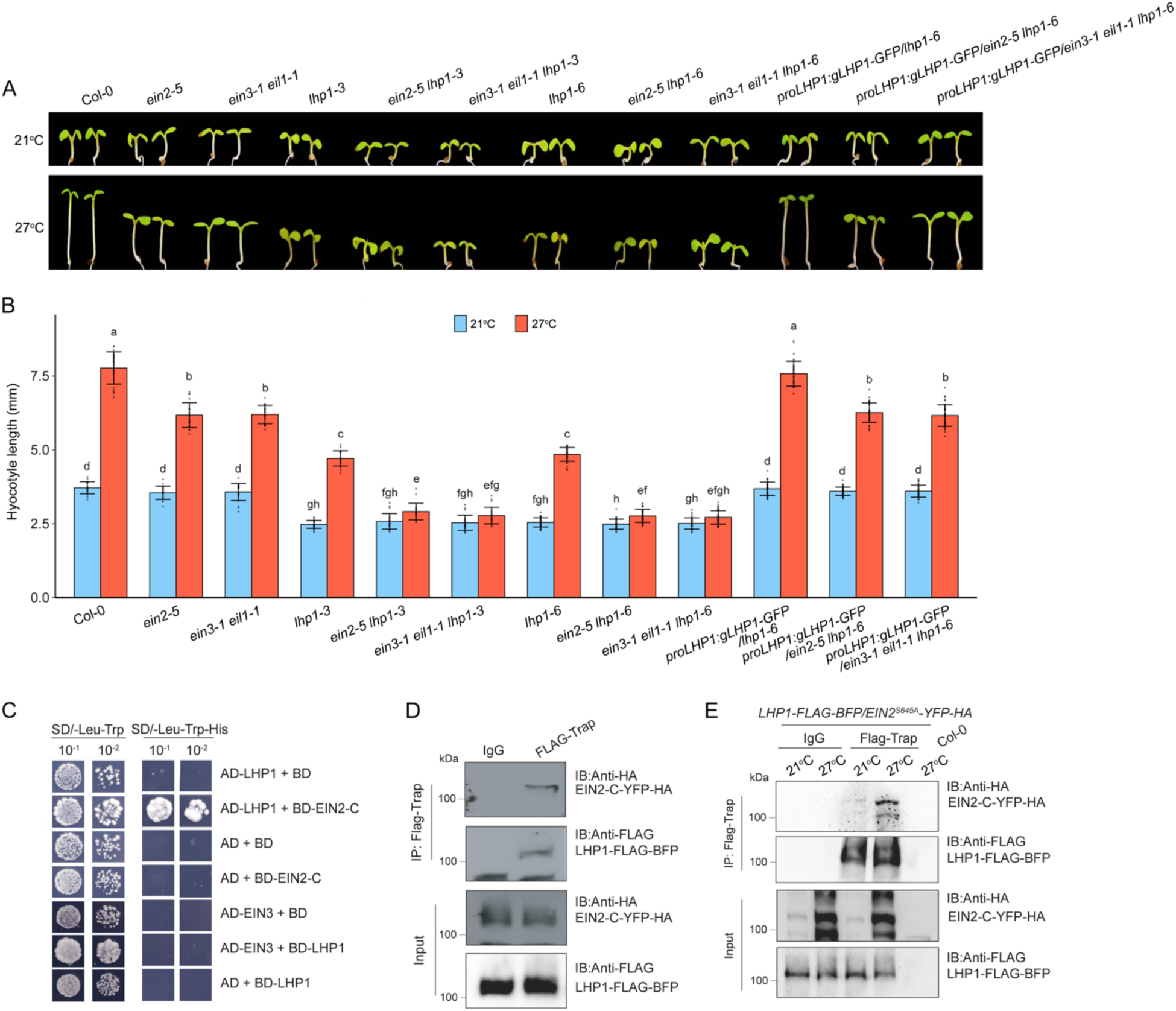
EIN2 associates with LHP1 to regulate warm ambient temperature response. (**A**) Representative images of seedlings of indicated plants grown on MS medium at either 21°C or 27°C. (**B**) Measurements of hypocotyl lengths of indicated plants in (**A**). Values are means ± SD of at least 30 seedlings. Different letters represent significant differences between each genotype calculated by a One-way ANOVA test followed by Tukey’s HSD test with *P* ≤ 0.05. (**C**) Yeast two-hybrid assays to detect the interaction between LHP1 and EIN2-C or EIN3. BD: GAL4 DNA binding domain; AD: GAL4 activation domain. LHP1 was fused to AD and used as bait proteins to test the interaction between EIN2-C and LHP1; AD-EIN3 was used as bait proteins to test the interaction between EIN3 and LHP1. Left panels: Yeasts grown on two-dropout medium (SD/-Leu-Trp) served as a loading control; Right panels: Yeast grown on selective three-dropout medium as experiment group (SD/-Leu-His-Trp). (**D**) Pull-down assays of EIN2-C (EIN2-C-YFP-HA) with LHP1 (LHP1-FLAG-BFP). The total proteins from the tobacco leaves co-transformed with *EIN2-C-YFP-HA* and *LHP1-FLAG-BFP* were applied to the immunoprecipitation by Flag-Trap magnetic agarose (DYKDDDDK Fab-Trap) by using LHP1-FLAG-BFP as bait. The pull-down products were detected in immunoblots against anti-HA antibody or anti-FLAG antibody. IgG-coated magnetic beads were used as negative control. (**E**) *In vivo* co-immunoprecipitation assay to examine the interaction between EIN2-C and LHP1. Five-day-old *LHP1-FLAG-BFP/EIN2^S645A^-YFP-HA* transgenic green seedlings treated with or without 4 hours of 27°C were subject to immunoprecipitation with Flag-Trap magnetic agarose (DYKDDDDK Fab-Trap), IgG magnetic beads as negative control. Anti-HA antibody or anti-FLAG antibody were used to detect IP products. IB, immunoblotting; IP, immunoprecipitation.

To understand the molecular basis of the synergistic function of EIN2, EIN3, and LHP1, we performed Y2H assays and *in vitro* pull-down assays and found that EIN2-C rather than EIN3 could interact with LHP1 (Fig. 1C and 1D). To further confirm the interaction between LHP1 and EIN2-C *in vivo* under the warm ambient temperature condition, we performed a co-immunoprecipitation (Co-IP) assay in the *LHP1-FLAG-BFP*/*EIN2^S645A^-YFP-HA* plants with or without four hours of 27°C treatment (Fig. 1E). An increased interaction between EIN2-C and LHP1 was detected when *35S::LHP1-BFP-FLAG/ EIN2^S645A^-YFP-HA* transgenic plants were treated with 27°C for four hours (Fig. 1E). These results provide an additional piece of biochemical evidence that LHP1 and EIN2 function collaboratively in responses to warm ambient temperature.

To further investigate the collective function of EIN2 and LHP1 in warm ambient temperature response, we performed mRNA-seq of *ein2-5lhp1-3* and compared the warm ambient temperature activated differentially expressed genes (DEGs) in *ein2-5lhp1-3* with those in *ein2-5* and *lhp1-3* single mutants and Col-0 (Fig. S2D). We found that 205 of DEGs were both EIN2- and LHP1-dependent as their log_2_FC value is the lowest in *ein2-5lhp1-3* than that in *ein2-5,* in *lhp1-3*, and in Col-0 (Fig. 2A). Gene ontology (GO) analysis showed that biological processes of response to heat, response to stimulus, and cellular response to stimulus were significantly enriched (Fig. S2E). We also noted that numbers of well-studied growth promoting and stress responsive genes were in this category (Fig. 2B).

**Figure 2.**
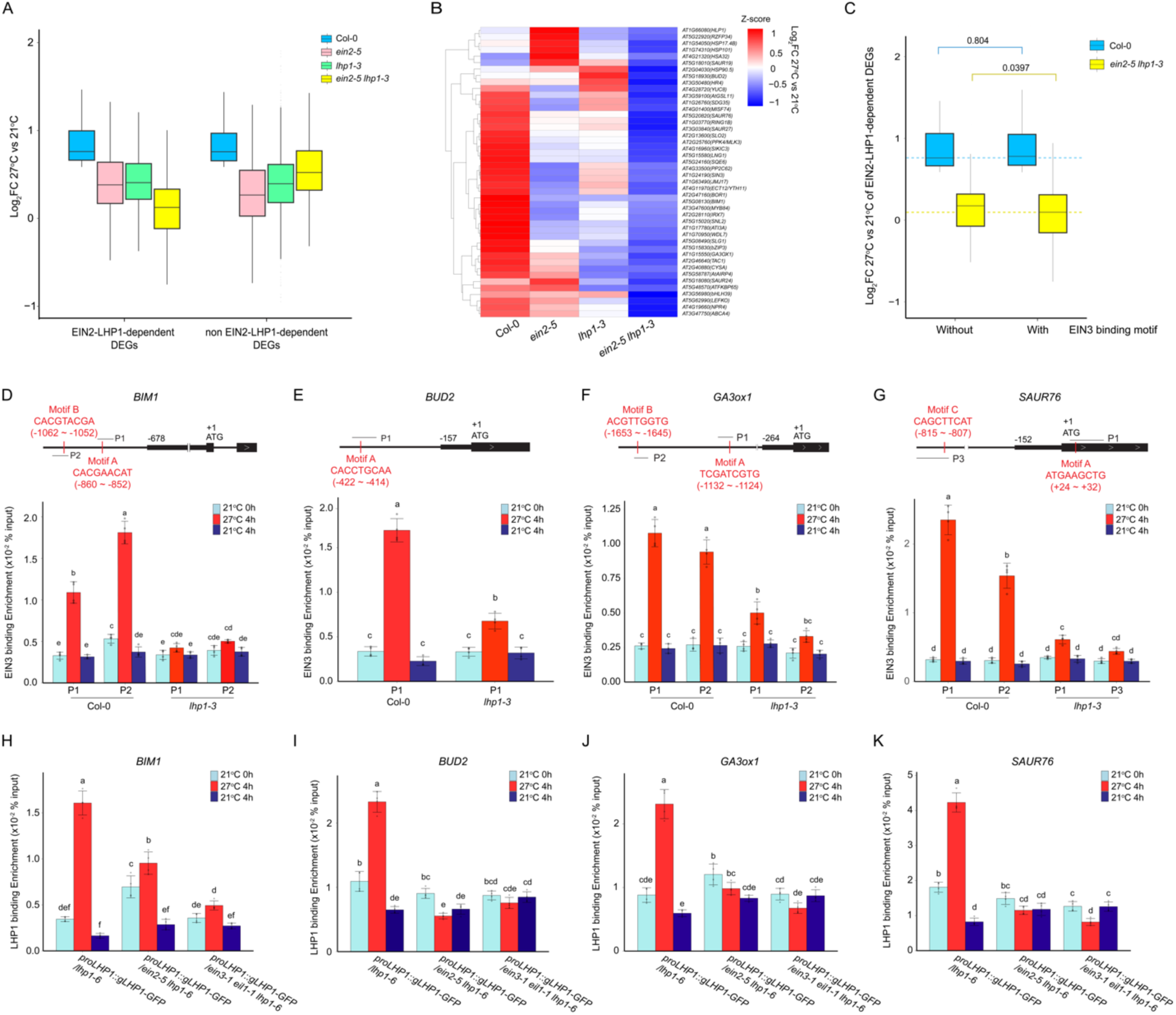
EIN2, EIN3, and LHP1 collectively regulate warm ambient temperature response at transcription level. (**A**) Box plots to compare the expression of EIN2-LHP1-dependent warm ambient temperature-induced DEGs versus those genes that are EIN2-LHP1 independent in Col-0, *ein2-5, lhp1-3,* and *ein2-5lhp1-3* plants. (**B**) Heatmap to show the transcriptional responsiveness to warm ambient temperature of key EIN2-LHP1 co-dependent DEGs in Col-0 and in the indicated plants. Z-score transformed log_2_FC value of each gene was used to plot the heatmap. (**C**) Box plots to compare the reduction of transcriptional activation in the EIN2-LHP1 co-dependent DEGs with EIN3 binding motif versus without EIN3 binding motif in Col-0 and *ein2-5lhp1-3* plants. *P* values were determined by a two-tailed *t* test. (**D**-**G**) ChIP-qPCR assay to detect EIN3 enrichment at EIN3 binding motifs in the target genes. The top panels are the diagrams to show the locations of ChIP-qPCR primers in the promoter regions of each target gene. Red vertical lines indicate EIN3 binding motifs. Black bracelets indicate the location of ChIP-qPCR amplicons. Chromatins isolated from five-day-old seedlings collected at 21°C before treatments, 27°C treatments for 4h, and 21°C control treatments for 4h were immunoprecipitated with anti-EIN3. (**H**-**I**) ChIP-qPCR assays to examine the enrichment of LHP1 on the target genes in the indicated genetic backgrounds. Five-day-old seedlings were collected at 21°C before treatments, 27°C treatments for 4h, and 21°C control treatments for 4h and subject to ChIP assay with anti-GFP. For (**D**-**I**), data represent the mean ± SD of at least three replicates and the original percentage of input values were plotted as dots. The statistically significant differences of EIN3 or LHP1 binding in different conditions and genetic backgrounds was calculated by a One-way ANOVA test followed by Tukey’s HSD test with *P* ≤ 0.05.

The genetic interaction between EIN3 and LHP1 and the phenotypic similarity between *ein3-1eil1-1lhp1-3* and *ein2-5lhp1-3* prompted us to investigate how EIN3 is involved in EIN2-LHP1-dependent warm ambient temperature induced transcriptional activation. We searched for the EIN3 binding motifs in the upstream 2000 bp (−2kb) from the start codon (ATG) in 205 of EIN2-LHP1-dependent DEGs using *de novo* EIN3 binding motif (A/T)(T/C)G(A/C/T)A(T/C/G)(C/G)T(T/G)^13^. 124/205 of EIN2-LHP1-dependent ambient temperature activated DEGs have at least one putative EIN3 binding motif (Fig. 2C). Notably, the reduction in transcriptional activation of 124 genes with EIN3 binding motif(s) was more significant than that of the genes without EIN3 binding motif in *ein2-5lhp1-3* (Fig. 2C). Our RT-qPCR assay of four representative target genes, *BIM1*, *BUD2*, *GA3ox1*, and *SAUR76* ^14, 15, 16, 17^ in different genetic backgrounds further confirmed that EIN3 is required for their transcriptional activation in response to warm ambient temperature (Fig. S3 and Fig. S4A-4D). Importantly, the decrease in warm ambient temperature-induced transcriptional activation resulting from *EIN3/EIL1* mutations was amplified in the *ein3-1eil1-1lhp1-3* triple mutants, which is similar to that observed in the *ein2-5lhp1-3* double mutant (Fig. 2C and Fig. S3). This implies that EIN3 plays a role in the EIN2-LHP1-dependent warm ambient temperature response.

Considering that EIN3 binding motifs have been identified *in silico* within the promoter regions of EIN2-LHP1-dependent DEGs, and EIN3 is required for their transcription activation in response to warm ambient temperature (Fig. 2C, Fig. S3, and Fig. S4A-4D), we examined EIN3 binding to those target genes in response to warm ambient temperature. ChIP-qPCR assay results showed that EIN3 binding was highly enriched when Col-0 seedlings were subject to warm ambient temperature treatment (Fig. S4A-4H and Fig. 2D-2G). Moreover, the elevated EIN3 binding by warm ambient temperature was significantly reduced in *lhp1-3* mutant, although there is no significant difference in the endogenous EIN3 levels between Col-0 and *lhp1-3* (Fig. 2D-2G and Fig. S4I).

As LHP enhances EIN3’s function in warm ambient temperature response, and LHP1 can associate with chromatin, we therefore analyzed published LHP1 ChIP-seq dataset^18^. The LHP1 binding enrichment was detected in our target genes and more importantly, there is a noteworthy preference for LHP1 binding in EIN2-LHP1-dependent DEGs compared to the entire Arabidopsis genome (Fig. S4J-4N). Subsequent ChIP-qPCR over selected target genes in *proLHP1::gLHP1-GFP/lhp1-6* transgenic plants showed that the binding affinity of LHP1 was enhanced by warm ambient temperature (Fig. 2H-2K). Additional ChIP-qPCR assays in *proLHP1::gLHP1-GFP/ein2-5lhp1-6* and *proLHP1::gLHP1-GFP/ein3-1eil1-1lhp1-6* demonstrated that warm ambient temperature induced LHP1 binding was largely compromised by the mutations of *EIN2* and *EIN3/EIL1* (Fig. 2H-2K). Taken together, these data provide compelling evidence that EIN2, EIN3, and LHP1 collectively function in the same complex on a subset of genes that are responsible for warm ambient temperature response. Additionally, there is a mutual regulatory interaction between EIN3 and LHP1 in their binding activities.

LHP1 is considered as a component of both Polycomb Repressive Complex 1(PRC1) and PRC2, and known to recognize H3K27me3 histone tails^19, 20, 21^. We therefore examined H3K27me3 levels over those EIN2-LHP1-dependent DEGs using publicly available H3K27me3 ChIP-seq data^22^. Intriguingly, H3K27me3 histone marks were highly enriched over those DEGs (Fig. 3A), which is in contrast with the canonical role of H3K27me3 as a transcriptional repression mark^23, 24^. Many studies have shown that the H3K27me3 marked chromatin regions can co-exist with active histone modifications such as H3K4me3 to establish a bivalent epigenetic state for rapid gene regulation in response to various internal or external signals^25, 26, 27^. Therefore, we examined H3K4me3 over EIN2-LHP1-dependent DEGs using published ChIP-seq data (Fig. 3B)^28^. 42 of EIN2-LHP1-dependent genes are characterized as bivalent genes, and 25 of these bivalent genes have EIN3 binding motifs (Fig. 3C and Fig. S5A-5D). More importantly, most of the 25 genes have been reported to play essential functions in plant growth and stress response, indicating that EIN2, EIN3, and LHP1 target a key subset of bivalent genes in response to high ambient temperature (Fig. S5E) ^14, 15, 16, 17, 29, 30, 31, 32, 33, 34, 35, 36, 37, 38, 39^.

**Figure 3.**
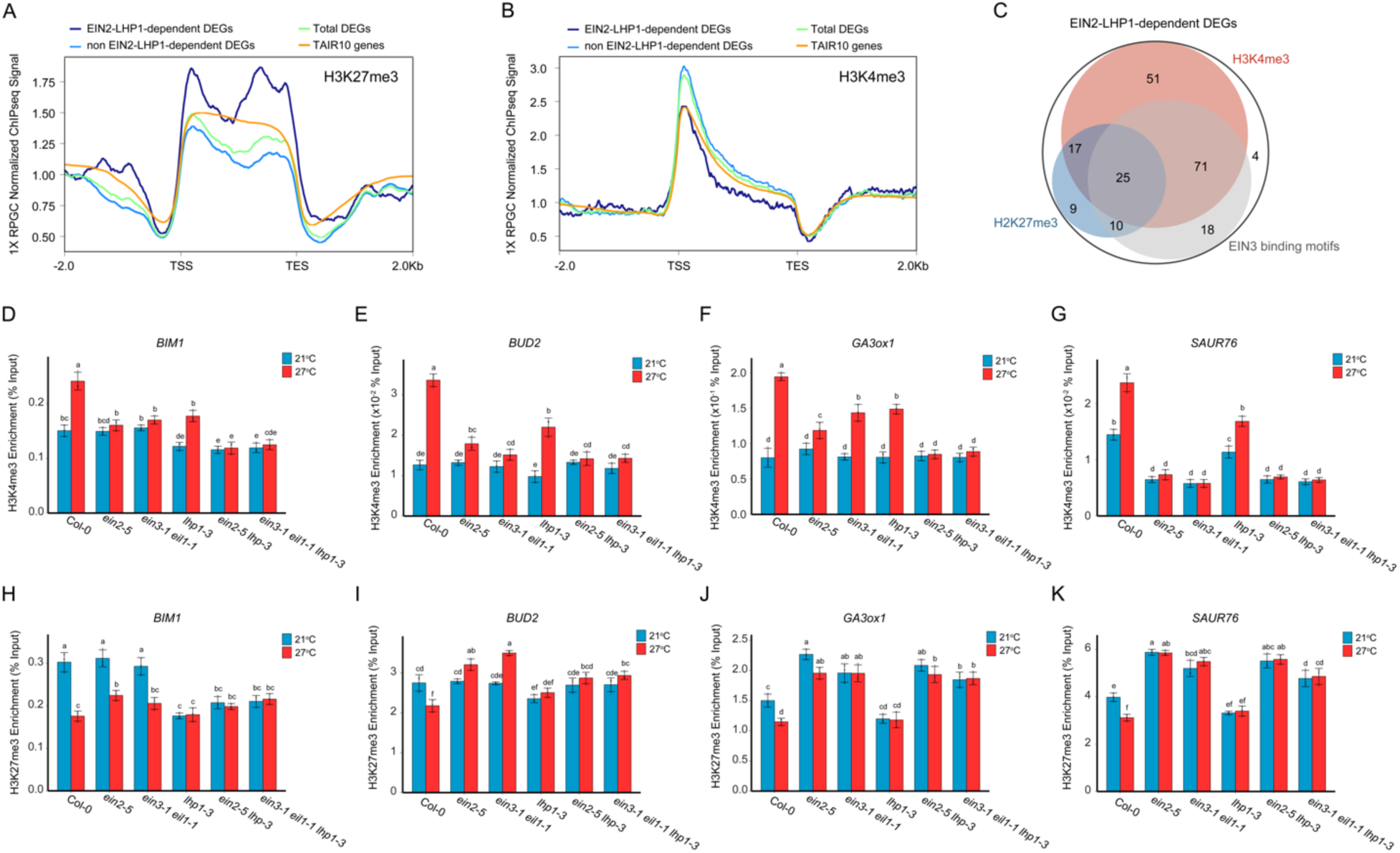
EIN2, EIN3, and LHP1 coregulate bivalent chromatin for warm ambient temperature induced transcriptional activation. (**A** and **B**) Profile plots of RPGC normalized H3K27me3 (**A**) and H3K4me3 (**B**) ChIP-seq signal over EIN2-LHP1 co-dependent warm ambient temperature induced DEGs, non EIN2-LHP1 co-dependent warm ambient temperature induced DEGs, total warm ambient temperature induced DEGs, and total TAIR10 annotated genes. TSS, transcription start site; TES, transcription end site. (**C**) Venn diagram to compare the H3K27me3 marked DEGs, H3K4me3 marked DEGs, and DEGs identified with EIN3 binding motifs in EIN2-LHP1 co-dependent warm ambient temperature induced DEGs. (**D**-**K**) ChIP-qPCR analyses to evaluate the enrichment of H3K4me3 (**D**-**G**) and H3K27me3 (**H**-**K**) in Col-0, *ein2-5, ein3-1eil1-1, lhp1-3, ein2-5lhp1-3,* and *ein3-1eil1-1lhp1-3* five-day-old green seedlings treated with 4 hours of 27°C or kept in 21°C for four hours of selected genes. Individual data point of the methylated histone enrichment relative to input is plotted as a dot. Different letters indicate statistically significant differences (*P* ≤0.05) between each experiment group calculated by a One-way ANOVA test followed by Tukey’s HSD test.

To assess the dynamics of H3K4me3 and H3K27me3 in response to warm ambient temperature in those bivalence genes, we performed ChIP-qPCR assay in the selected target genes. H3K4me3 levels were elevated by warm ambient temperature treatment in Col-0 (Fig. 3D-3G). However, the elevations were significantly reduced in *ein2-5, ein3-1eil1-1*, and *lhp1-3* mutants. Notably, the elevation was nearly undetectable in both *ein2-5lhp1-3* and *ein3-1eil1-1lhp1-3* mutants (Fig. 3D-3G). In contrast, H3K27me3 levels showed a slight reduction or remained similar to that in normal temperature in Col-0 in response to warm ambient temperature (Fig. 3H-3K). The reduction of H3K27me3 levels was abolished in *ein2-5* and *ein3-1eil1-1* related mutants. Together, these results demonstrate that EIN2, EIN3, and LHP1 collectively regulate the warm ambient temperature-induced H3K4me3 elevation to counteract repressive mark H3K27me3, facilitating gene transcriptional activation in bivalent chromatin.

Mutations in INO80 leads to reduction in warm ambient temperature response and INO80 is involved in the regulation of H3K4me3 deposition for active transcription for thermomorphogenesis^40^. Therefore, we decided to evaluate the involvement of INO80 in the ethylene mediated warm ambient temperature response. We generated *ein2-5ino80-5* double mutant and *ein3-1eil1-1ino80-5* triple mutants, and warm ambient temperature response phenotypic assay showed that the partial irresponsiveness of *ein2-5, ein3-1eil1-1,* and *ino80-5* was significantly enhanced in the *ein2-5ino80-5* and the *ein3-1eil1-1ino80-5* mutants (Fig. 4A and 4B). We also generated *lhp1-3ino80-5* double mutant and observed that the *lhp1-3ino80-5* double mutant showed a similar defective warm ambient temperature response as observed in the *ein2-5ino80-5* and *ein3-1eil1-1ino80-5* mutants (Fig. 4A and 4B). Moreover, the adult plants of *ein2-5ino80-5*, *ein3-1eil1-1ino80-5,* and *lhp1-3ino80-5* exhibited a severe developmental defect when compared to *lhp1-3* and *ino80-5* single mutants and Col-0 (Fig. S6A). Additionally, the reduced warm ambient temperature response in *ein2-5ino80-5, ein3-1eil1-1ino80-5* and *lhp1-3ino80-5* mutants was rescued by overexpressing *INO80,* further confirming that the mutation in *INO80* leads to additive defects for warm ambient temperature response in *ein2-5, ein3-1eil1-1,* and *lhp1-3* mutants, respectively (Fig. 4A and 4B, and Fig. S6B). Overall, these results demonstrate the genetic interactions among EIN2, EIN3, LHP1, and INO80 in response to warm ambient temperature.

**Figure 4.**
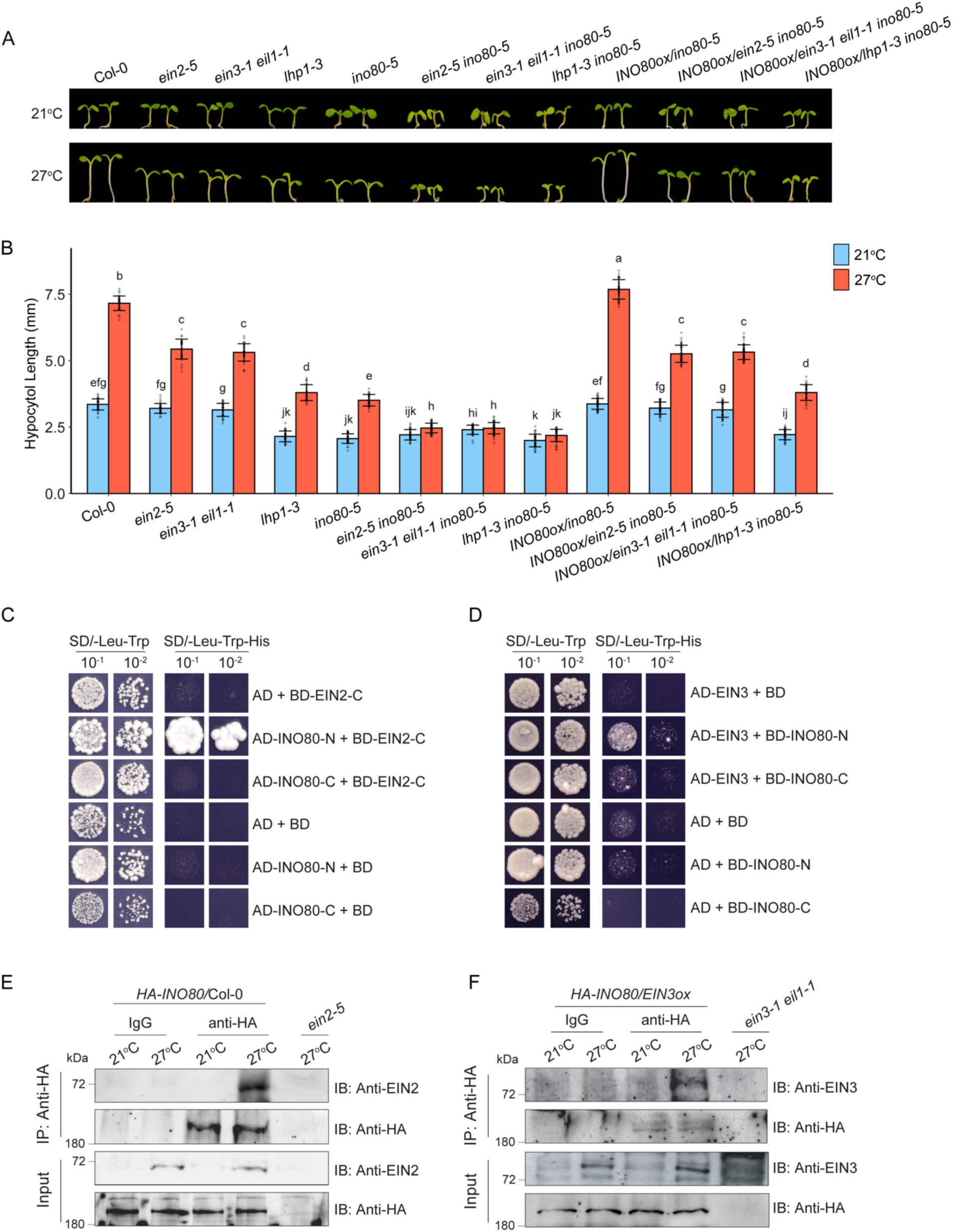
INO80 functions with EIN2 and EIN3 by protein-protein interaction for warm ambient temperature response. (**A**) Photographs of representative five-day-old green seedlings of Col-0, *ein2-5, ein3-1eil1-1, lhp1-3, ino80-5, ein2-5ino80-5, ein3-1eil1-1ino80-5, lhp1-3ino80-5,* and complementation lines by *35S::INO80* grown on MS medium at 21°C and 27°C. (**B**) Measurements of hypocotyl lengths from indicated plants in (**A**). Each value is means ± SD of at least 30 seedlings. Different letters indicate the statistically significant differences between different genotypes and treatments with *P* ≤ 0.05 calculated by a One-way ANOVA test and followed by Tukey’s HSD test for multiple comparisons. (**C** and **D**) Y2H assay to examine the interaction of INO80 (truncated INO80-N and INO80-C) with EIN2 (**C**) and EIN3 (**D**). AD, GAL4 activation domain; BD, GAL4 DNA binding domain. Left panels: Yeasts grown on two-dropout medium as loading control. Right panels: Yeasts grown on selective three-dropout medium. (**E** and **F**) *In vivo* co-immunoprecipitation assays of INO80 with EIN2 (**E**) and INO80 with EIN3 (**F**). The total protein extracts from five-day-old seedlings of *35S::HA-INO80*/Col-0 (**E**) or crossed *35S::HA-INO80/EIN3ox* (**F**) transgenic plants treated with 4 h at 21°C or 27°C were immunoprecipitated with anti-HA. The immunoprecipitated proteins were detected by immunoblotting using anti-HA, anti-EIN2, and anti-EIN3, respectively. The input serves as the loading control. IB, immunoblotting; IP, immunoprecipitation.

We then explored the potential physical interaction between INO80 and EIN2 or EIN3. We performed Y2H assays and found that the N terminus of INO80 (INO80-N) can interact with both EIN2-C and EIN3 *in vitro* (Fig. 4C and 4D, and Fig. S6C). Further *in vivo* coimmunoprecipitation assay showed that INO80 interacted with EIN2-C or EIN3 when seedlings were subjected to four hours of warm ambient temperature treatment (Fig. 4E, 4F and Fig. S6D). To further evaluate the functional interaction of INO80 with EIN2, EIN3, and LHP1, we first examined the target gene expression in *INO80* related mutants in response to warm ambient temperature. The result showed that the warm ambient temperature-induced activation of target genes was reduced in the *ino80-5* mutant, and this reduction was further enhanced in *ein2-5ino80-5, ein3-1eil1-1ino80-5,* and *lhp1-3ino80-5* mutants (Fig. S7A-S7D). We then examined H3K4me3 and H3K27me3 levels in the target genes in Col-0 and *ino80-5*. ChIP-qPCR results showed that H3K4me3 levels were decreased in *ino80-5* both with and without warm ambient temperature treatments, compared to those in Col-0 (Fig. 5A). In contrast, no significant differences in H3K27me3 levels were detected between in *ino80-5* and in Col-0 under normal condition. However, the warm ambient temperature-induced reduction detected in Col-0 was diminished in *ino80-5* (Fig. 5B).

**Figure 5.**
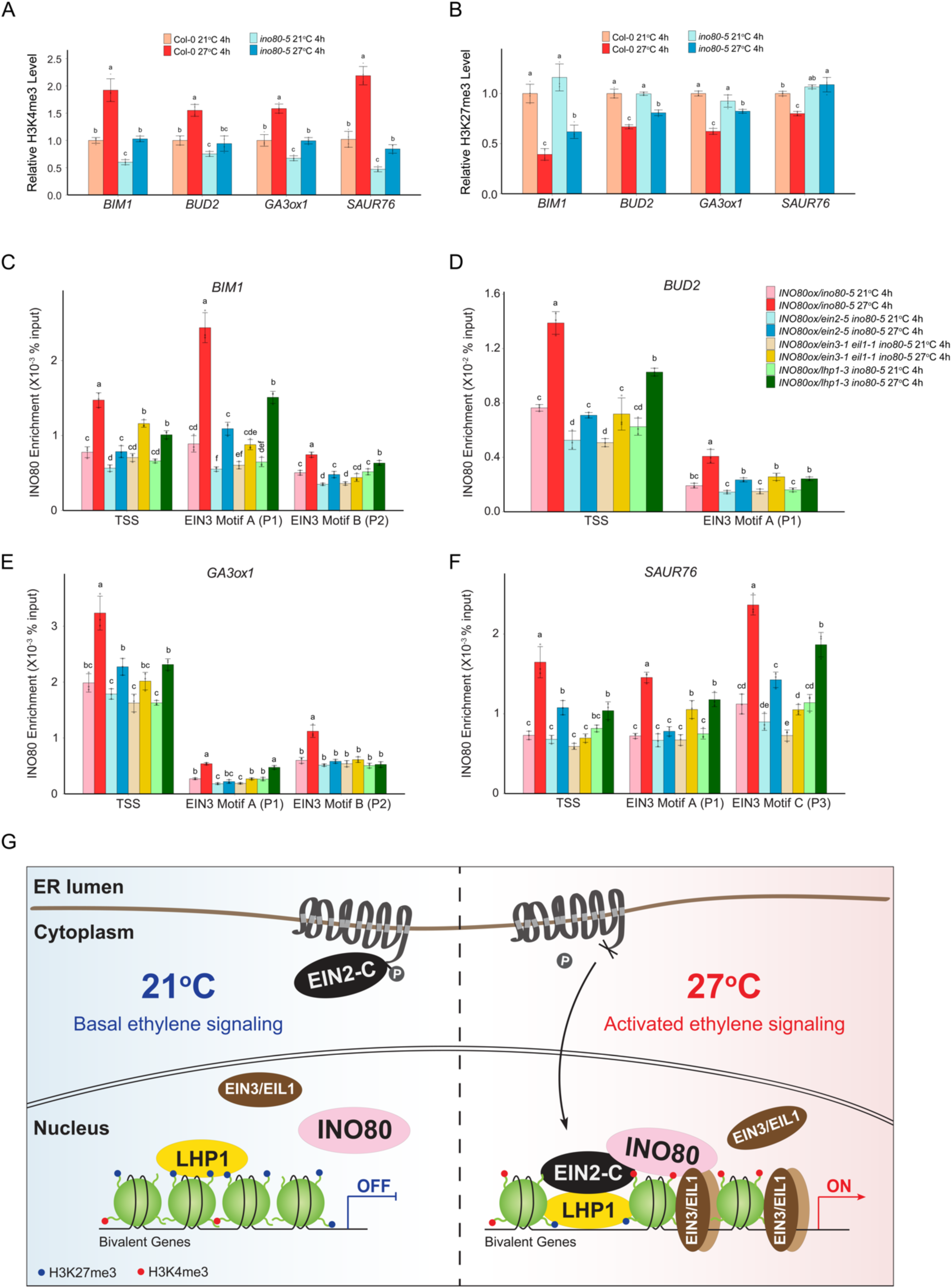
EIN2 and EIN3 recruit INO80 to bivalent chromatins to elevate H3K4me3 enrichment in response to warm ambient temperature. (**A** and **B**) ChIP-qPCR analyses of H3K4me3 (**A**) and H3K27me3 (**B**) enrichments on the target genes in Col-0 and *ino80-5* mutant in response to warm ambient temperature. Col-0 and *ino80-5* seedlings treated for four hours under 27°C or kept in 21°C for four hours were used for the ChIP assay. (**C**-**F**) ChIP-qPCR assays to examine the INO80 binding in *INO80ox/ino80-5, INO80ox/ein2-5ino80-5, INO80ox/ein3-1eil1-1ino80-5, INO80ox/lhp1-3ino80-5* in response to warm ambient temperature. Primers targeting TSS regions are the same primers used for H3K4me3 and H3K27me3 ChIP-qPCR for each gene. Primers targeting EIN3 binding motifs are the same primers used for EIN3 ChIP-qPCR for each gene in Fig. 2H-2K. All data represent means ± SD of three replicates and individual data point was plotted as dot. Different letters represent significant differences with *P* < 0.05 in the one-way ANOVA test followed by Tukey’s HSD test. (**G**) A schematic model to illustrate how LHP1 and INO80 cooperate with EIN2 and EIN3 to regulate bivalent chromatin for rapid transcriptional activation in response to warm ambient temperature. In normal temperature condition, LHP1 binds to H3K27me3 heavily labeled bivalent genes and those genes are transcribed at basal level. On the other hand, warm ambient temperature condition triggers the activation of ethylene signaling, resulting in EIN2 C-terminal cleavage and EIN3 protein accumulation. LHP1 directs EIN2-C to bivalent chromatin, where EIN2-C recruits INO80 and EIN3 to a subset of genes with bivalent marks to elevate H3K4me3 enrichment, counteracting the repressive effects of H3K27me3 and ultimately activating gene expression.

Since INO80 interacts with EIN3 and is associated with H3K4me3, we tested the INO80 binding at both EIN3 binding motifs and TSS regions of target genes in the ChIP-qPCR assays. Remarkably, INO80 displayed an increased binding affinity to the target genes under warm ambient temperature while INO80 protein levels remained unchanged (Fig. 5C-5F and Fig. S6D). Moreover, INO80 binding was significant reduced when *EIN2, EIN3,* or *LHP1* was mutated (Fig. 5C-5F). Accordingly, expressing INO80 alone is not sufficient to increase H3K4me3 levels in the absence of EIN2, EIN3, and LHP1 in response to warm ambient temperature; further ChIP-qPCR assay showed that INO80 had no effects on H3K37me3 levels (Fig. S7E-S7L), supporting our hypothesis that EIN2 and EIN3 recruit INO80 to LHP1-EIN2 and EIN3 co-targeted chromatin regions to regulate H3K4me3. Finally, we explored whether INO80 regulates EIN3 binding given the fact that activated bivalent chromatin regions have been shown to gain accessibility for transcription factor binding to activate gene expression^41, 42, 43^. In the absence of INO80, the binding of EIN3 to selected target genes was decreased, although there is no difference in its protein level in Col-0 and in *ino80-5* (Fig. S7M-S7Q). Altogether, our results demonstrate that EIN2 and EIN3 recruit INO80 to specific bivalent targets to elevate H3K4me3 enrichment for the activation of gene expression in response to warm ambient temperature.

In this study, we reveal that LHP1 and INO80 cooperate with ethylene signaling for warm ambient temperature response by activating specific bivalent genes. First of all, both our genetic and molecular results clearly showed that ethylene signaling and LHP1 share a similar regulation in warm ambient temperature (Fig. 1 and Fig. S1). More importantly, our genetic results showed the synergetic regulation of LHP1 with EIN2 and EIN3 in warm ambient temperature response (Fig. 1 and 2). As LHP1 is a component of PRC complexes that is potentially involved in the regulation of H3K27me3 and functions as a H3K27me3 reader^19, 21^, our new discovery established a link between ethylene signaling and H3K27me3 in warm ambient temperature response. Interestingly, the genes that are activated by both LHP1 and ethylene signaling in response to warm ambient temperature have a relative higher level of H3K27me3 enrichment (Fig. 3A), which is in contrary with the classical function of H3K27m3 as a transcriptional repression mark^44^. Our epigenetic profiling analyses showed that the subset of warm ambient temperature responsive DEGs whose transcriptional activation requires EIN2, EIN3, and LHP1 are marked with both H3K4me3 and H3K27me3 (Fig. 3A-3C). More importantly, H3K4me3 is elevated by warm ambient temperature to counteract H3K27me3 for transcriptional activation, which requires all EIN2, EIN3, and LHP1 (Fig. 3). Furthermore, we identified INO80 as the key factor that is recruited by EIN2-LHP1-EIN3 complex to promote H3K4me3 deposition for transcription activation (Fig. 4 and 5). Overall, our findings highlight the importance of the ethylene signaling pathway and its coordination with bivalent chromatin regulation for gene transcription in response to warm ambient temperature (Fig. 5G).

## Materials and methods

### Plant materials and growth conditions

All *Arabidopsis thaliana* plants used in this study were in the Col-0 ecotype background and grown at continuous temperatures of 21°C or 27°C in the growth chamber set with same light intensity condition and an 8-h-light/16-h-dark short-day (SD) light cycle. The phenotypic assays in response to warm ambient temperature were performed according to previous publication^45^. In brief, approximately 50 seeds of each genotype collected at the similar time were surface sterilized, plated on the same Murashige & Skoog (MS) medium supplemented with 1% sucrose, and placed at 4°C in the dark for three days for stratification. After germination was induced at 21 °C for 24 hours, seedlings were either shifted to 27°C SD conditions or kept at 21°C SD as control till day 5. Their hypocotyl lengths were measured using Fiji ImageJ software^46^. The mutants *ein2-5, ein3-1ein1-1, lhp1-3, lhp1-6, ino80-5,* and *pif4-2* were used in this study. Higher order mutants were generated by genetic crossing. *EIN2, EIN3, LHP1*, and *INO80* related transgenic plants were derived from previous publications^12, 40, 47, 48^, except *35S::LHP1-FLAG-BFP/*Col-0 (generated in this study). To obtain *proLHP1::gLHP1-GFP/lhp1-6, proLHP1::gLHP1-GFP/ein2-5lhp1-6,* and *proLHP1::gLHP1-GFP/ein3-1eil1-1lhp1-6* complementation lines, *proLHP1::gLHP1-GFP/lhp1-6 FRI* was backcrossed to Col-0, *ein2-5,* and *ein3-1eil1-1* to remove *FRI,* respectively; *lhp1-6*, *ein2-5lhp1-6,* and *ein3-1eil1-1lhp1-6* were also generated from the same crossed population by segregation. Homozygous plants of both *proLHP1::gLHP1-GFP* transgene and related gene null mutations with comparable *gLHP1-GFP* expression levels were used for following analyses. The *35S::HA-INO80/ein2-5ino80-5*, *35S::HA-INO80/ein3-1eil1-1ino80-5,* and *35S::HA-INO80/lhp1-3ino80-5* lines were generated by genetic crossing *35S::HA-INO80/ino80-5* with *ein2-5, ein3-1eil1-1,* and *lhp1-3*, respectively.

### Plant protein extraction and western blot assays

To assess warm ambient temperature response at the protein level, seedlings in each indicated genetic background after warm ambient temperature treatment were frozen and homogenized in liquid nitrogen. Total protein was extracted with extraction buffer (100 mM Tris-HCl, pH 7.5, 100 mM NaCl, 5 mM EDTA, 5 mM DTT, 10 mM β-mercaptoethanol, 1% SDS, 1 mM PMSF, and 1X protease inhibitors from Thermo Fisher), and the plant debris was removed by centrifugation. The extracted proteins in the supernatant were further denatured at 85 °C for 5 min after mixing with 2x Laemmli sample buffer and subject to SDS-PAGE. The protein levels were detected by immunoblot analysis using the primary antibodies: anti-HA antibody (Biolegend, #901503, dilution 1:5000), anti-FLAG antibody (CST, #14793, dilution 1:2000), anti-EIN2 antibody (raised in-house, dilution 1:2000)^47^, anti-EIN3 antibody (raised in-house, dilution 1:2000)^48^, anti-GFP antibody (Invitrogen, A11122, dilution 1:2000), and anti-PIF4 (Agrisera, AS16 3955, dilution 1:2000) and corresponding secondary antibodies: anti-mouse (Bio-Rad, #105001G, dilution 1:10000), anti-rabbit (Bio-Rad, #1706515, dilution 1:10000), and anti-goat (Agrisera, AS09 605, dilution 1:5000). HRP activity was detected using enhanced chemiluminescence (ECL; GE Healthcare) according to the manufacturer’s instructions with either ChemiDoc Imaging System (Bio-Rad) or conventional X-ray films.

### Plant RNA extraction and gene expression analysis

To assess transcriptional response to warm ambient temperature treatment, five-day-old green seedlings in each indicated genetic background grown in 21°C SD were shifted to warm ambient temperature (27°C) from ZT4 for four hours till ZT8 in SD condition or kept at 21°C till ZT8. Plants samples were collected at 21°C 0h (ZT4 21°C, before treatment), 27°C 4h (ZT8 27°C, warm ambient temperature treatment), and 21°C 4h (ZT8 21°C, control treatment) and immediately snap-frozen in liquid nitrogen. Total RNA was extracted after frozen seedlings were homogenized in liquid nitrogen using the RNeasy Plant Kit (Qiagen) with DNase I digestion and clean up (Qiagen, RNeasy Mini Kit (250)). First-strand cDNA was synthesized using Invitrogen Superscript^TM^ III First-Strand cDNA Synthesis Kit. cDNAs and qPCR primers were combined with 2X SYBR master mix from Thermo Fisher for qPCR. PCR reactions were performed in triplicates on a Roche 96 Thermal cycler. *ACTIN2* was used as endogenous control for normalization. Three independent biological replicates were performed with reproducible results. Primers used in this study are listed in Supplementary Table 1.

### mRNA sequencing and data processing

RNA was extracted from five-day-old green seedlings with or without four hours of 27°C treatment of each genotype using the RNeasy Plant Kit (Qiagen). For mRNA sequencing (mRNA-seq) library preparations, libraries were generated using the NEBNext® Poly(A) mRNA Magnetic Isolation Module (E7490) and the NEBNext® Ultra™ II RNA Library Prep Kit for Illumina® (E7770), with 1μg RNA as input, following the manufacturer’s instructions. Indexed libraries were sequenced on the platform of HiSeq2000 (Illumina). Paired-end raw fastq data were evaluated with FastQC and low-quality reads were removed with Trim Galore (Babraham Institute). The trimmed and filtered reads were then mapped to the Arabidopsis reference genome (TAIR10) with STAR^49^. Mapped reads were counted by featureCounts (Subread 2.0.1) and differentially regulated genes were identified using R package DESeq2^50^ with a p-adj< 0.05 and | log2(fold change) | > 0.585 (|fold change | > 1.5). The boxplots, violin plots, and dot plots were generated by R package ggplot2 and the heatmaps of log2FC were generated by R package pheatmap. GO enrichment analysis for DEGs was performed by agriGO v2^51^. Trimmed and aligned read counts are listed in Supplementary Table 2.

### Yeast Two-Hybrid Assay

The yeast two-hybrid assay was performed according to previous publications using the ProQuest Two-Hybrid System (Invitrogen) following the manufacturer’s instructions^52, 53^. Briefly, the bait and prey plasmids (pAD-LHP1, pBD-LHP1, pAD-EIN3, pBD-EIN2-C, pAD-INO80-N, pBD-INO80-N, pAD-INO80-C, and pBD-INO80-C) were co-transformed into the yeast strain AH109, following the experiment design. For INO80 related Y2H assays, INO80 full length protein was truncated to INO80-N (1-500aa) and INO80-C (501-1540aa) for optimal expression in yeast according to literature^40^. The positive transformants were selected and grown on SD/-Leu-Trp (two-dropout) medium. 3-amino-1,2,4-triazole (3-AT) was supplemented to repress self-activation. Growth on SD/-Leu-Trp-His (three-dropout) medium supplemented with appropriate concentrations of 3-AT indicates the physical interaction between corresponding proteins.

### Transient expression in *Nicotiana benthamiana* leaves

To co-express EIN2-C-YFP-HA and LHP1-FLAG-BFP fusion proteins in *N. benthamiana* for pull-down assays, 20 ml overnight cultures of *Agrobacterium tumefaciens* (strain GV3101) in LB medium carrying the *35S::EIN2-C-YFP-HA pEarleyGate101* and *35S::LHP1-FLAG-BFP pCambia1300* binary vectors respectively were pelleted by centrifuge at 2000 g at room temperature for 15 min and then resuspended in infiltration buffer (10 mM MgCl_2_, 10 mM MES, pH 5.7, and 100 μM acetosyringone, AS). Paired resuspended Agrobacterium cultures were combined and then mixed with an equal volume of p19 agrobacteria resuspension to reach a final OD600 of 0.8 for each resuspension before the infiltration at the abaxial side of *N. benthamiana* leaves using a needleless syringe. After growth for 48-72 hours under dim light condition, infected tobacco leaves were collected and snap frozen in LN2 for further protein analysis.

### Co-immunoprecipitation

Co-immunoprecipitation (Co-IP) assays between proteins of interest were performed according to previous publications^54, 55^. In brief, soluble total proteins were extracted from fine ground plant tissues in two volumes of co-IP buffer (50 mM Tris-Cl, pH 8, 150 mM NaCl, 1 mM EDTA, 0.1% Triton X-100, 1 mM PMSF, and 1x protease inhibitor cocktail) at 4°C for 15 min with gentle rocking. The lysates were cleared by centrifugation at 4°C (10 min, 5000 g). For HA-INO80 fusion protein related co-IP experiments, the anti-HA antibody (Biolegend, #901503) was prebound to the equilibrated IgG coated Dynabeads (Thermo Fisher) for three hours with gentle rotation at 4°C. For LHP1-FLAG-BFP fusion protein related co-IP assays, DYKDDDDK Fab-Trap™ agarose beads (ChromoTek) were used to immunoprecipitate LHP1 protein complexes, following the same procedure. A 10% input aliquot for each co-IP experiment was taken from the cleared supernatant before the rest of the supernatant was added to magnetic beads and incubated at 4°C with gentle rocking overnight. IgG coated Dynabeads were incubated with protein extracts as negative control for each IP experiment. Magnetic beads with IP products were precipitated magnetically using a DynaMagnetic rack (Thermo Fisher) and then washed five times with 1 mL of co-IP buffer. Immunoprecipitated proteins were then released from the magnetic beads using 2x Laemmli sample buffer by heating at 85°C for 8 min and subject to SDS-PAGE.

### ChIP sequencing data analysis

Publicly available ChIP sequencing (ChIP-seq) datasets of LHP1, H3K27me3, and H3K4me3 were obtained from the Gene Expression Omnibus (GEO) database^18, 22, 28^. Raw sequencing data were subject to quality control by FastQC. Low quality reads were removed with Trim Galore (Babraham Institute) and then mapped to the Arabidopsis genome with Bowtie 2 (2.4.2)^56^. Duplicate reads were removed using SAMtools and broad peak calling for methylated H3 was performed using MACS2 with ‘--broad’ option^57^. Histone modification peaks were assigned to corresponding genes if a peak overlaps with the proximal region of a gene, including 1.5 kb upstream of TSS and 1.5 kb downstream of TES, by ChIPseeker^58^. ChIP-seq coverage tracks shown in the genome browser IGV were generated using the bamCoverage function in deeptools3.0.2^59^ normalized by reads per genomic content (1x normalization, 1X RPGC) and bin size = 1. The ChIP-seq signals from 2kb upstream TSS to 2kb downstream TES of indicated warm ambient temperature related DEG groups were calculated with bamCompare and computeMatrix and plotted with plotProfile function in deepTools.

### ChIP-qPCR assays

Chromatin immunoprecipitation was performed according to previous publications^60, 61^. In brief, five-day-old green seedlings shifted to 27°C for four hours or kept at 21°C were harvested at the same time (ZT8, SD) and crosslinked in 1% formaldehyde. For EIIN3 and LHP1 related ChIP-qPCR assays, plant samples prior to the four-hour treatment were also collected at 21°C ZT4 (21°C, 0h) as EIIN3 and LHP1 are reported to be regulated by light cycle and circadian clock^62, 63^. Nuclei were isolated and the chromatin was sheared by Bioruptor to approximate 200-300 bp fragments. Anti-GFP (Invitrogen, A11122, for LHP1 binding), anti-EIN3, anti-H3K27me3 (Active Motif, 61017), anti-H3K4me3 (Active Motif, 39016), or anti-HA (Biolegend, #901503, for INO80 binding) antibodies together with Dynabeads were added to the sonicated chromatin followed by incubation overnight to precipitate protein bound DNA fragments. The amount of immunoprecipitated DNA was quantified by qPCR and normalized to input DNA. Three independent biological replicates were performed for each line and condition with reproducible results. Primers used for ChIP-qPCR are listed in Supplemental Table 1.

## Acknowledgements

We thank C. Dean for sharing *proLHP1::gLHP1-GFP/lhp1-6 FRI* seeds. We thank D. Jiang for sharing *35S::HA-INO80/ino80-5* and *ino80-5* seeds. We thank J. Kim and S. Sung for suggestions on the experiment design. This work was supported by grants from the National Institute of Health to HQ (NIH-2R01 GM115879).

## Contributions

Z.S. and H.Q. conceived the project and designed the experiments. Z.S. performed most of the experiments and data analyses. Y.B. performed PIF4 immunoblots. Z.S. and H.Q. wrote the manuscript. E.H. advised on the experiment design and the manuscript. All authors read and approved the final manuscript.

## Data and materials availability

Further information and requests for all unique materials generated in this study can be directed to and will be fulfilled by the corresponding author Hong Qiao. The high-throughput sequencing data generated in this study have been deposited in the Gene Expression Omnibus (GEO) database (accession no. GSE256453).

